# Microvasculature-on-a-Chip: Bridging the interstitial blood-lymph interface via mechanobiological stimuli

**DOI:** 10.1101/2021.04.08.438936

**Authors:** Barbara Bachmann, Sarah Spitz, Christian Jordan, Patrick Schuller, Heinz D. Wanzenböck, Bahram Haddadi, Michael Harasek, Heinz Redl, Wolfgang Holnthoner, Peter Ertl

**Affiliations:** Faculty of Technical Chemistry, Institute of Applied Synthetic Chemistry and Institute of Chemical Technologies & Analytics, TU Wien, 1060 Vienna, Austria; Ludwig Boltzmann Institute for Experimental and Clinical Traumatology in the AUVA Research Centre, 1200 Vienna, Austria; Competence Center MechanoBiology, 1200 Vienna, Austria; Austrian Cluster for Tissue Regeneration, 1200 Vienna, Austria; Institute of Chemical, Environmental and Bioscience Engineering, Thermal Process Engineering and Simulation, TU Wien, 1060 Vienna, Austria; Institute of Solid State Electronics, TU Wien, Vienna, 1040, Austria

**Keywords:** Vascularization, Sprouting, Lymphatic System, Organ-on-a-Chip, Mechanobiology

## Abstract

After decades of simply being referred to as the body’s sewage system, the lymphatic system has recently been recognized as a key player in numerous physiological and pathological processes. As an essential site of immune cell interactions, the lymphatic system is a potential target for next-generation drug delivery approaches in treatments for cancer, infections, and inflammatory diseases. However, the lack of cell-based assays capable of recapitulating the required biological complexity combined with unreliable *in vivo* animal models currently hamper scientific progress in lymph-targeted drug delivery. To gain more in-depth insight into the blood-lymph interface, we established an advanced chip-based microvascular model to study mechanical stimulation’s importance on lymphatic sprout formation. Our microvascular model’s key feature is the co-cultivation of spatially separated 3D blood and lymphatic vessels under controlled, unidirectional interstitial fluid flow while allowing signaling molecule exchange similar to the *in vivo* situation. We demonstrate that our microphysiological model recreates biomimetic interstitial fluid flow, mimicking the route of fluid *in vivo*, where shear stress within blood vessels pushes fluid into the interstitial space, which is subsequently transported to the nearby lymphatic capillaries. Results of our cell culture optimization study clearly show an increased vessel sprouting number, length, and morphological characteristics under dynamic cultivation conditions and physiological relevant mechanobiological stimulation. For the first time, a microvascular on-chip system incorporating microcapillaries of both blood and lymphatic origin *in vitro* recapitulates the interstitial blood-lymph interface.

## INTRODUCTION

The lymphatic system absorbs, transports, and recirculates excess interstitial fluid via initial lymphatic capillaries, collecting lymphatic vessels, lymph nodes, and the thoracic duct. Even though this process is essential for physiologic fluid homeostasis, the historical view of the lymphatic system as a simple fluid conduit has needed reconsideration in the last decade following discoveries of its vital function in immune surveillance,^1^ inflammation,^2^ and cancer metastasis.^3^ Aside from its role as a significant component of the immune system, playing a crucial part in autoimmune disease and infection,^4,5^ the lymphatic system is a preferential site for cancer metastasis,^6^ facilitated by decreased fluid stress and lymph nodes as holding reservoirs.^7^ With the growing understanding of lymphatic physiology and disease, the interest in targeting the lymphatic system for drug delivery for potency enhancement of anti-inflammatory drugs, vaccines, or chemotherapeutics has risen accordingly. During physiological fluid homeostasis, interstitial fluid flow pushes excess fluid, molecules, and cells towards the initial lymphatic capillaries. Within the initial lymphatics, lymph formation and molecule uptake occur via passive transport through endothelial cell junctions. The fate of these molecules within interstitial tissue predominantly depends on their size. Small molecules preferentially reenter into blood capillaries, while lymphatic capillaries drain larger molecules such as proteins, pathogens, or immune cells.^8^ Current methods in lymphatic research to study these transport phenomena, however, rely heavily on animal models.^9^ While valuable to determine the influence of specific genes on lymphatic function *in vivo*, these models often lack the spatiotemporal resolution to track single-cell and molecule movement under controlled and reproducible measurement conditions.

Traditional *in vitro* models of vascular biology, in contrast, often involve endothelial monolayers, transwell cultures, or pseudo-capillary structures on hydrogels and simplify biology to a point where effective translation to *in vivo* phenomena becomes questionable. Alternatively, organ-on-a-chip technology provides the opportunity to engineer microphysiological systems within an *in vivo*-like microenvironment optimally tailored to the individual research question, overcoming traditional *in vitro* models’ limitations.^10–12^ In particular, microfluidic vascular cell culture systems demonstrate successful vascularization to form interconnected, perfusable three-dimensional (3D) vascular networks between two medium channels.^13^ To provide a brief overview, Table 1 lists recent publications on 3D microvascular systems and highlights how microfluidic approaches are preferentially used to study the vascular microenvironment.

**Table 1:** Recent reports and applications of microfluidics for 3D vascular biology. IF = interstitial flow.

| VASCULATURE TYPE | CULTIVATION SETUP | MICROFLUIDIC | APPLICATION | REF |
| --- | --- | --- | --- | --- |
| BLOOD | Vascular network | Yes | Vasculogenesis and growth factor gradients | 14 |
| BLOOD | Sprouting | Yes | IF magnitude and direction | 15 |
| BLOOD | Vascular network | Yes | Endothelial paracellular and transcellular permeability | 16 |
| BLOOD | Vascular network | Yes | Measurement of trans-capillary solute flux | 17 |
| LYMPH | Sprouting | Yes | Influence of IF on sprouting lymphangiogenesis | 18 |
| LYMPH | LEC in hydrogel | Yes | Contribution of lymphatic endothelium on colorectal cancer cell growth | 19 |
| BLOOD/LYMPH | Single Vessel | Yes | Anti-angiogenic drug screening | 20 |
| BLOOD/LYMPH | Vascular network | No | Integration of blood and lymph capillaries in tissue-engineered constructs | 21 |
| BLOOD/LYMPH | 2D separated by a membrane | Yes | Blood- and lymphatic-vessel permeability assay | 22 |

Among these, self-assembled and autologously formed vascular networks in hydrogel-laden microfluidic channels have proven ideal for studying fundamental biological processes, diseases, and exposure to drugs or toxic compounds.^16,23,24^ The success of vasculature-on-chip systems is inherently linked to the unique capability to adjust physiological flow conditions,^15^ shear forces,^25^ nutrient gradients,^14^ and biomechanical cues.^11,26^ Additionally, the characteristic design flexibility of microfluidic technology has led to the generation of a variety of customized cultivation chamber geometries capable of precisely controlling cell-cell, cell-matrix, and cell-molecule interactions.^11,12,27^

Vessel formation *in vivo* occurs either via vascularization from single cells or via sprouting angiogenesis from preexisting vasculature. Sprouting angiogenesis relies on cues from the surrounding tissue signaling nutrient demand or excess fluid accumulation. In other words, endothelial cells can sense mechanobiological stimuli arising from fluid flow irregularities in the interstitial tissue and respond by forming new vessels to regain fluid homeostasis. Similarly, both blood^15^ and lymphatic^18^ endothelial cells *in vitro* react sensitively to flow velocity and direction, dependent on their physiologic function. Despite recent advances in mimicking blood vasculature, models of the lymphatic system remain scarce. To date, only one study demonstrates the cultivation of 3D blood and lymphatic vessels on a microfluidic platform,^20^ even though such systems could investigate immune cell chemotaxis or lymphedema progression.^28^ As an example, mechanobiological stimulation via directional interstitial fluid flow increases tumor cell entry into lymphatic capillaries and stimulates lymphatic sprouting.^29,18^ Emulating the mechanobiological environment regarding matrix elasticity,^30^ fluid flow,^31^ or nutrient gradients^32–34^ is crucial for developing reliable and accurate *in vitro* systems. We have previously shown that mechanobiological control over shear stress and growth factor gradients allows manipulation of vascular network formation and nanoparticle uptake by endothelial cells.^14,31^.

The lessons learned from mimicking the cardiovascular endothelium in microphysiological systems provide a feasible opportunity to establish, for the first time, an interstitial blood-lymph interface featuring 3D microcapillaries of both blood and lymphatic origin.^35^ Building on our previous knowledge, we now developed a microfluidic system containing two adjacent and locally separated hydrogel-embedded microvascular systems to study the impact of interstitial fluid flow regimes and mechanobiological stimulation on lymphatic vessel formation. A hydrostatic pressure-driven fluid flow inside the microvasculature-on-chip system accomplishes mechanobiological stimulation resulting in a unidirectional, interstitial-like fluid flow. Within our optimized blood-lymph interface, primary microvascular blood and lymphatic endothelial cells are co-cultured in the presence of adipose-derived stem cells under static and dynamic conditions. Consequently, our study sets out to investigate the impact of directional interstitial fluid flow regimes on vascularization and sprouting angiogenesis and lymphangiogenesis, leading to the formation of a microvascular blood-lymph interface.

## MATERIALS AND METHODS

### Microfabrication

Microfluidic devices were manufactured using soft-lithography techniques to generate hybrid poly-dimethylsiloxane (PDMS) / glass microdevices. Photolithographic molds were produced using Ordyl SY 300 dry-film photoresist (Elga Europe) on silicon wafers to generate 100 µm high replicas of the microdevice design. Microfluidic structures were attained by liquid PDMS (Sylgard® 184, Dow Corning) casting and subsequent polymerization at 80°C for 90 minutes. After generating hydrogel inlet ports and medium reservoirs using 1 mm and 6 mm biopsy punches, microfluidic chambers were bonded to clean glass slides using oxygen plasma (30 seconds, 80% power). Three microfluidic chambers are featured on one object slide-sized microchip for facilitated replica cultivation as visible in a microfluidic device rendered in Figure 1A. These vascularization chambers feature two individually loadable hydrogel chambers separated by an array of pillar structures 100 µm in diameter. Before cell loading, microfluidic devices were sterilized using 70 % Ethanol and dried for at least 24h to ensure the restoration of hydrophobic properties.

**Figure 1:**
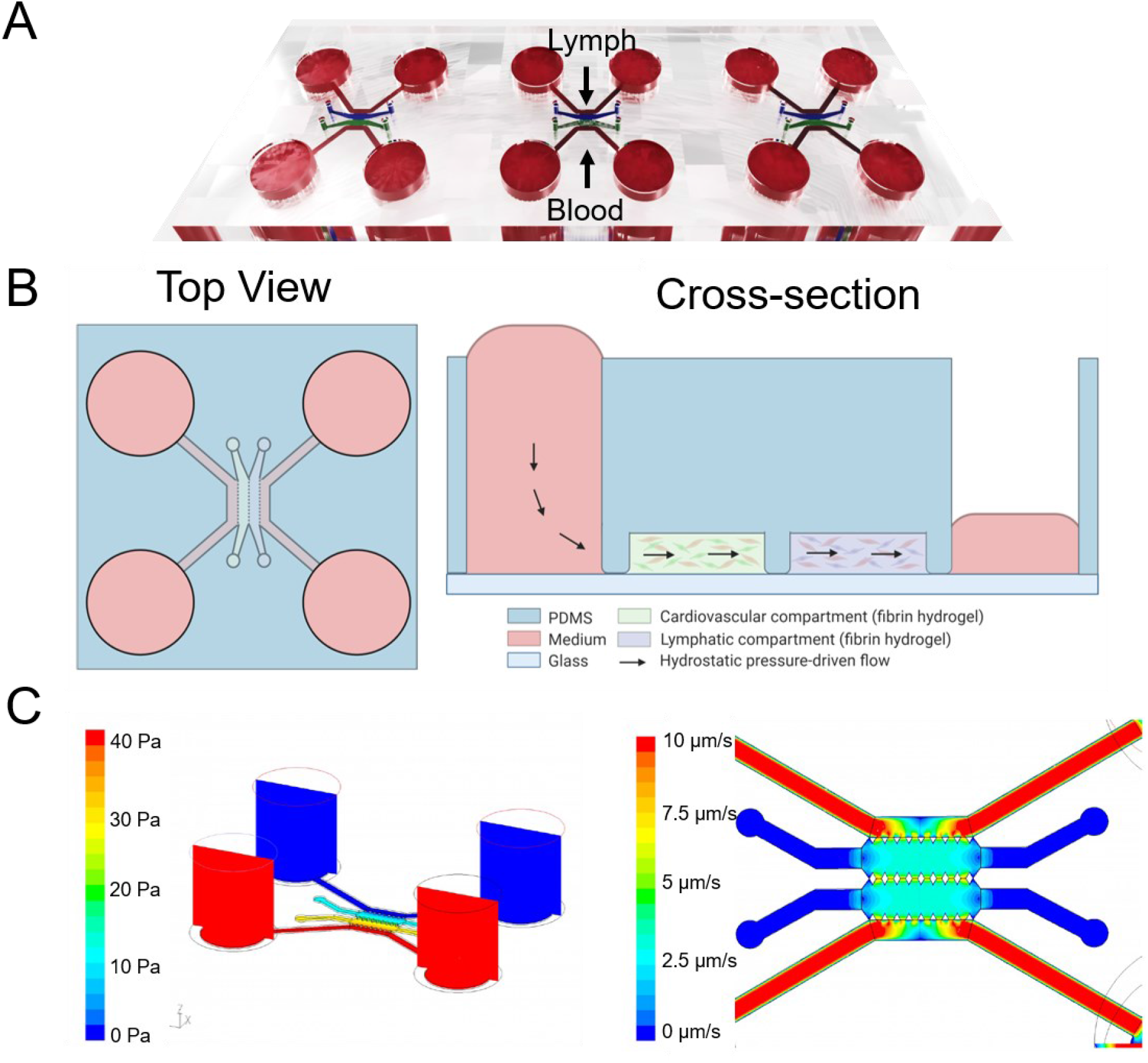
A) Render of Microvessel-on-chip device housing three separately addressable cell cultivation chambers on one object-slide sized microchip. (B) Schematics of a top view and cross-section of one cell cultivation chamber designed for side-by-side cultivation of blood and lymphatic endothelial vasculature stimulated via unidirectional hydrostatic pressure-driven flow. (C) CFD simulation of hydrostatic pressure-driven flow in microvessel-on-chip device controlled by medium reservoir fluid level. Pressure-driven flow within the hydrogel area represents interstitial flow-like velocities ranging between 1 – 3 µm/s.

### Computational fluid dynamic (CFD) simulations

A commercial general-purpose CFD code (Ansys Fluent 6.3.26) was utilized for simulating fluid flow and concentrations within the device. The fluid flow in a microchip can be characterized as laminar (Re << 1), for the fluid-wall interaction, a no-slip boundary condition has been applied. Given the low species concentrations and assuming Newtonian behavior, the fluid properties of water were used in the model (density 998 kg/m^3^, dynamic viscosity 1 mPa.s). Hydrogels were modeled using an isotropic porous medium approach with a fixed porosity and a constant viscous resistance (0.99 and 6.67 ^10-12^/m, respectively). Diffusion coefficients were used from literature data.^33^ A specified ambient temperature (25 °C) and constant atmospheric pressure (1 atm = 101325 Pa) were set as reference conditions. At the inlet and outlet boundary zones of the flow geometry, constant velocity and/or pressure boundary conditions were derived from the experimental setups. For transient simulation runs, grid cell size and time steps were matched to meet the Courant stability criterion to ensure physically valid results. To speed up simulation run time, gravity forces were replaced by equivalent pressure on the boundaries for steady-state snapshot simulations. The simulation results (“virtual experiments”) can also be utilized to derive more generic correlations as a description of, e.g., a function describing the fluid flow rate depending on the filling level for a given chip geometry and physical parameter set. From multiple steady-state CFD calculations, flow fields, concentration gradients, and accumulation or depletion of reservoir fluids as well as dissolved species in defined fluid regions have been monitored and illustrated.

### Cell culture

Primary human umbilical vein endothelial cells (HUVEC) and adipose-derived stem cells (ASC) were isolated and cultivated as previously described^21,36^ after approval by the local ethics committees of Upper Austria and the AUVA and written donor consent. Telomerase-immortalized lymphatic endothelial cells (LEC-tert) were a gift from Marion Gröger (Medical University of Vienna), and primary microvascular blood endothelial cells (BEC), and lymphatic endothelial cells (LEC) were obtained from PromoCell. All cells were cultivated in a fully supplemented endothelial cell growth medium (EGM-2, Promocell) with 5 % fetal bovine serum (FBS), if not recommended otherwise by the supplier, and used for on-chip experiments before passage 10. HUVEC and LEC-tert were retrovirally transfected to express green fluorescent protein (GFP) or red fluorescent protein (mCherry) as previously published.^37^

### Microchip loading and induction of vascular network formation

For microchip loading of hydrogel-embedded cell suspensions, bovine fibrinogen (Sigma Aldrich) was resuspended in PBS^-/-^ at 6 mg/mL and dissolved by incubation at 37°C for three hours before sterile filtering. Cells were washed with PBS, detached from cell culture flasks, and resuspended in an FBS-free endothelial cell medium before the addition of bovine thrombin (Sigma Aldrich). Fibrinogen and cell/thrombin solutions were subsequently mixed and injected into the hydrogel region using 8 µL cell suspension/chamber to yield final concentrations of 3 mg/mL fibrinogen, 1 U/mL Thrombin, 5*10^6^ endothelial cells/mL, and 2.5*10^6^ ASC/mL. Fibrin hydrogels were polymerized for 10 minutes at room temperature in humidity chambers as previously described^38^ before the second hydrogel chamber was filled following either the procedure mentioned above or with cell-free fibrin hydrogel. Subsequently, hydrogels were polymerized for 20 minutes at 37°C before supplying cell culture medium via the fluid reservoirs. For lymphatic sprouting experiments, medium channels were coated with 1 µg/mL fibronectin for 1h before injecting 5 µL LEC suspension at 5*10^6^ cells/mL, and devices were tilted by 90° to ensure cell attachment to the hydrogel interface. Microvascular cultures were maintained in EGM-2 or a mixture of EGM-MV (PromoCell) and EGM-MV2 (Promocell), depending on cultured cell types, with daily medium changes for five days. Primary lymphatic endothelial cell cultures were supplied with vascular endothelial growth factor C (VEGF-C, CoaChrom Diagnostica) at 25 ng/mL, 50 ng/mL, or 100 ng/mL and stimulated with 100 ng/mL Phorbol 12-Myristate 13-Acetate (PMA) as described. Selected devices were perfused using 70 kDa TRITC-dextran (Sigma Aldrich) via hydrostatic pressure-driven flow.

### Immunofluorescent staining

For immunofluorescent staining, microvascular devices were washed twice in HBSS, fixed in 4 % paraformaldehyde (PFA) for 30 minutes, and permeabilized using 0.1 % Triton-X100 for 15 minutes. Specimens were blocked overnight in a blocking buffer of 2 % goat serum in HBSS, incubated with rabbit anti-VE-cadherin antibody (Abcam) for 72h, washed, and bound to goat anti-rabbit secondary antibody (Abcam) for 48h. Nuclei were stained using DAPI at 1 µg/mL, and F-actin fibers were visualized using a 1x phalloidin solution (Abcam). Images were obtained on an Olympus IX83 epifluorescent microscope using lasers at appropriate wavelengths.

### Image and data analysis

Image analysis of microvascular networks and lymphatic sprouting behavior was performed using Image J and AngioTool software.^39^ Data were visualized and analyzed using GraphPad Prism. Where applicable, outliers were excluded using a Grubbs Outlier test at α = 0.05, and statistical significance was determined at p-value < 0.05 using Student’s t-tests or two-way ANOVA.

## RESULTS AND DISCUSSION

### Microfluidic device supports spatially separated cultivation and generates interstitial flow

Directional flow conditions are vital for maintaining the blood-lymph interface *in vivo*, where blood endothelial cells sprout in flow direction, and lymphatic endothelial cells need to sprout in the opposite direction of the fluid flow.^15,18^ We integrated two separately loadable hydrogel compartments for blood and lymphatic vasculature to investigate whether unidirectional flow also guides endothelial cell sprouting inside a microfluidic biochip. These compartments are in close proximity and downstream from each other to allow reciprocal cell signaling and cell-to-cell interactions. In total, the microvasculature-on-chip device, depicted in Figure 1A, contains three individually addressable cell culture units (µLymph units) on a single device to enable twelve measurements in parallel using a microwell plate-sized chip holder. Each of the individual µLymph units, shown in Figure 1B, further consists of two cell culture chambers, two medium channels, and four medium reservoirs. Equidistant circular micropillars separate the individual hydrogel-laden chambers from each other and the respective medium channels ensuring (1) reproducibility of hydrogel injection, (2) individual nutrient and growth factor supply to each of the cultures, (3) reciprocal signaling between the cell types, and (4) establishment of a directional fluid flow across both hydrogel regions. Controlling fluid column height within the reservoirs enables a directional fluid flow from the reservoir across the adjacent hydrogel chambers containing the blood and lymphatic vasculatures. For instance, a fluid column difference of 4 mm between blood and lymphatic fluid reservoirs creates a pressure drop of 40 Pa across the hydrogel region, as confirmed by CFD simulation illustrated in Figure 1C. This pressure difference results in a mean unidirectional fluid flow across the hydrogel region of 2 µm/s, thus mimicking the pressure gradient in the interstitial capillary bed. Detailed CFD simulation data depicting column height and fluid flow velocities over four days of culture, shown in Figure S1, describe peak fluid flow velocities of 3 µm/s after the medium change and a decrease until column equilibration at day 4. These results correlate well with interstitial fluid flow velocities found *in vivo* ranging between 0.1 and 4 µm/s, depending on tissue type, location, and activity level.^40^ By simply adjusting reservoir fill height every 24 h, our microvasculature-on-chip device can maintain physiologic interstitial flow velocities between 1 – 3 µm/s for mechanobiological stimulation of endothelial cell behavior over extended cultivation periods (e.g., weeks).

### Lymphatic sprouting setup ideal for determination of flow-induced sprouting variations

Next, we investigated the suitability of our microvasculature-on-chip device to mimic the blood-lymph interface. Initially, a cell culture optimization study was performed in the presence and absence of interstitial fluid flow using blood endothelial cells (HUVEC) and lymphatic endothelial cells (LEC-tert) in co-culture with adipose-derived stem cells (ASC), as a replacement of vessel supporting cells such as pericytes.^21^ Figure 2A shows a schematic drawing of the dual vascularization setup (left) and the lymphatic sprouting setup (right). HUVEC and ASC are simultaneously co-seeded as a cell suspension in fibrin hydrogel within the left hydrogel chamber in both cultivation setups. The main difference between dual vascularization and lymphatic sprouting setups is that in the lymphatic sprouting configuration, LEC-tert are not mixed into the fibrin hydrogel but instead added via the exit channels to form an endothelial cell monolayer at the hydrogel interface by tilting the microfluidic device during cell attachment. Hence, LEC-tert need to sprout into the hydrogel to form vessels that reach and interact with the blood vasculature. Results of our cell culture optimization study, shown in Figure 2B and S2A, revealed a stable and reproducible formation of blood vasculature and endothelial sprouting within a five-day cultivation period. Additional perfusion studies using 70 kDa TRITC-dextran (see Figure S3) confirmed the functionality and structural integrity of the on-chip generated blood vascular networks. A direct comparison of the intertwined blood-lymphatic capillaries generated in the absence (static) and presence of fluid flow (dynamic) using the dual vascularization setup showed no discernible impact of interstitial flow conditions on vascularization behavior. In turn, the presence of unidirectional interstitial fluid flow positively impacted lymphatic endothelial sprouting as early as day 3 of culture. Mechanobiological stimulation initiated sprouting activity early on, leading to endothelial sprout elongation under dynamic growth conditions and pointing at the activation of sprouting behavior by shear stress perpendicular to the endothelial cell surface. White arrows in Figure 2B (right lower panel) further highlight lymphatic sprout formation at day 5 of culture, thus highlighting our microvessel-on-chip’s ability to achieve an *in vitro* blood-lymphatic capillary interface using both cultivation approaches.

**Figure 2:**
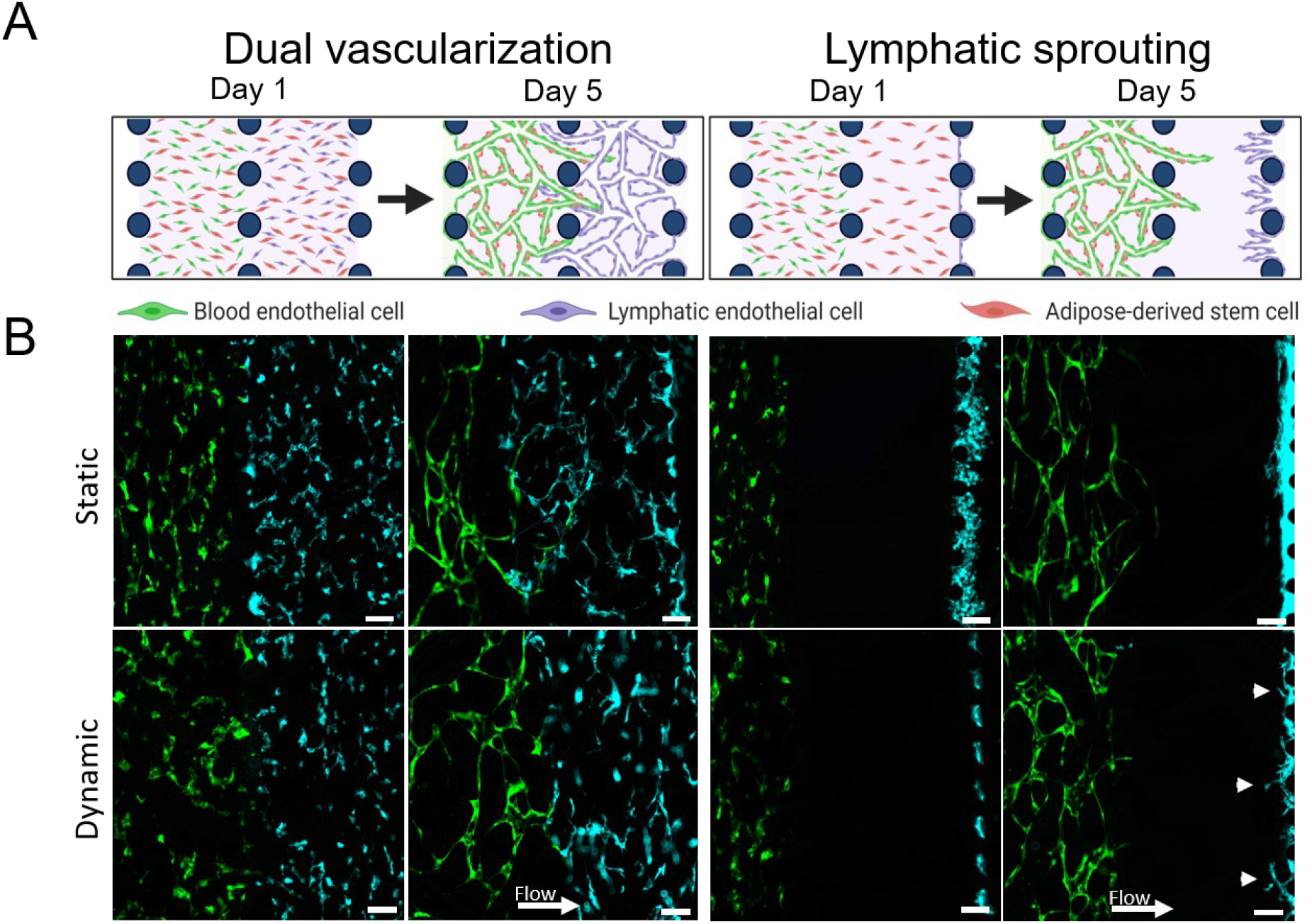
Cultivation strategies performed using the MechanoLymph device depicted as a sketch (A) and fluorescent endothelial cells (B). Blood endothelial cells (HUVEC-GFP) form vascular networks in co-culture with adipose-derived stem cells (ASC, not labeled) seeded within fibrin hydrogel within the left chamber of the device. Lymphatic endothelial cells (mCherry LEC-tert, shown in cyan) are seeded within the right compartment either as hydrogel-embedded culture (dual vascularization setup) or as cell monolayer adherent to the hydrogel at the medium interface (lymphatic sprouting setup) and form lymphatic vasculature via vasculogenesis or sprouting into the hydrogel within five days of culture. Dynamic cultivation has no discernible effect in the dual vascularization setup but enhances lymphatic sprouting. Scale Bar is 100 µm.

### Primary microvascular LEC respond to the cooperative effect of VEGF-C and PMA

In the next set of experiments, primary microvascular lymphatic endothelial cells (LEC) replaced LEC-tert as a model system to avoid any effects attributed to telomerase-mediated prolongation of cellular life span and to attain a more physiologically relevant chip-based lymphatic model. For optimal culture conditions, the impact of growth factor concentration on lymphatic sprouting was determined in exclusively static conditions using the lymphatic sprouting set up described above using increasing vascular endothelial growth factor C (VEGF-C)^41^ without or with additional stimulation via phorbol 12-myristate 13-acetate (PMA)^42^. The growth factor optimization study results in respective sprout number and sprout length changes, both key quality parameters, as shown in Figure 3. Interestingly, primary LECs highly rely on PMA stimulation to form lymphatic sprouts in static culture since very limited sprouting occurred only in 25 ng/mL VEGF-C non-PMA stimulated cultures. A combination of PMA (100 ng/mL) and VEGF-C (25 ng/mL) stimulation, however, resulted in consistent lymphatic endothelial sprout formation (see Figure 3A).

**Figure 3:**
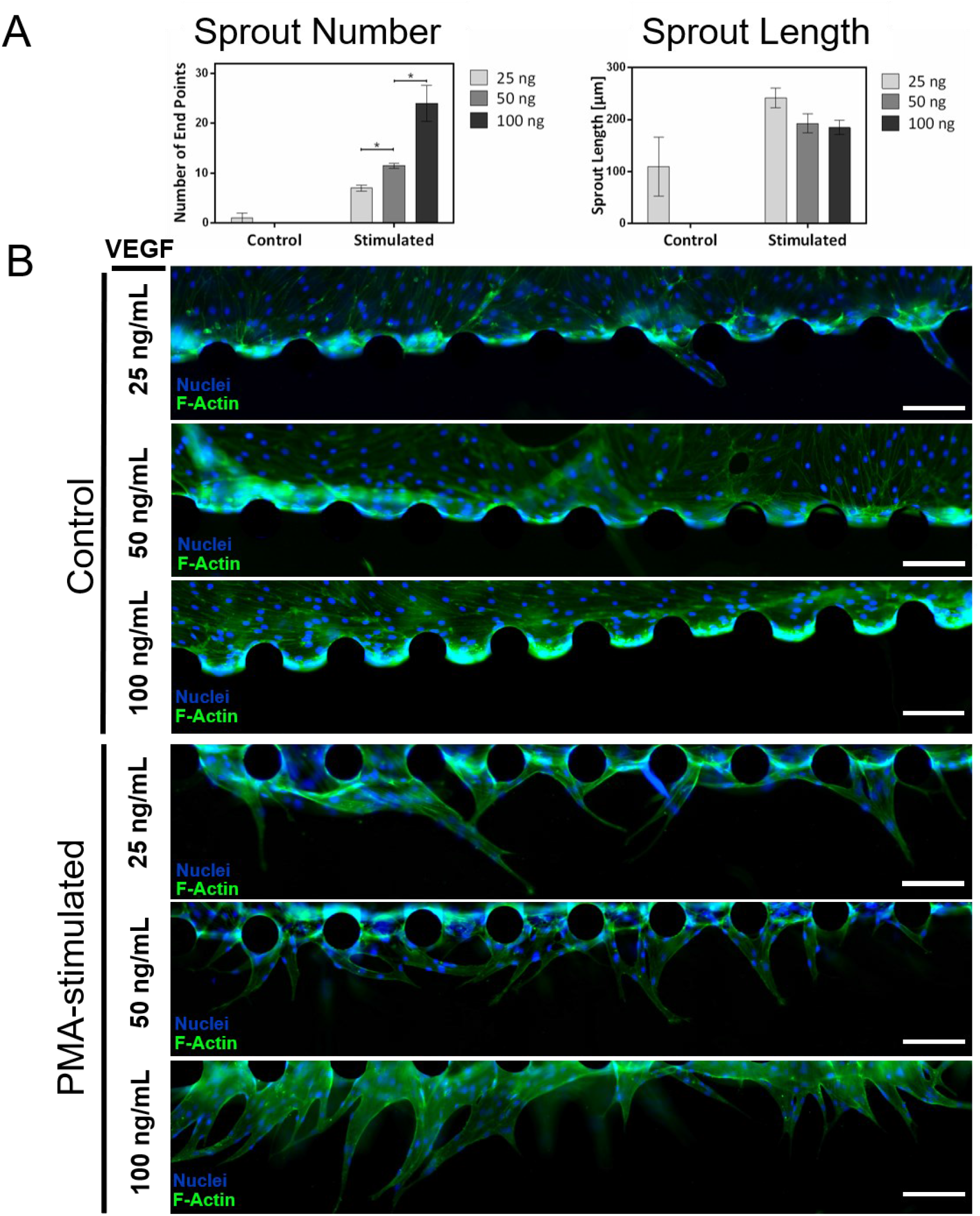
(A) Increase of lymphatic sprout number with VEGF-C concentration in culture medium and addition of PMA and differences in sprout length with VEGF-C concentration. (B) Epifluorescent images of LEC in culture with varied VEGF-C concentration and PMA stimulation. Stimulated LEC show increased sprouting behavior and morphological differences dependent on VEGF-C concentration. Scale Bar 200 µm. *p<0.05.

Moreover, when increasing the VEGF-C concentration from 25 to 50 and 100 ng/mL, the mean sprout number rose from 7 to 24 sprouts per device in the presence of 100 ng/mL PMA (see Figure 3A). It is important to note that a similar sprout length of approx. 200 µm (e.g., 242 µm, 193 µm, and 185 µm) was obtained independent of the applied VEGF-C concentration in PMA-stimulated cultures. Sprout morphology, however, deviated slightly dependent on VEGF-C stimulation. For instance, LEC treated with 25 ng/mL or 50 ng/mL formed fine capillaries often reconnecting to lumenized vessels after initial sprout growth and featuring single tip cell protrusions that extend into the cell-free hydrogel as visualized by nuclei and F-actin staining in Figure 3B. In contrast, cultures stimulated with 100 ng/mL VEGF-C emanated from the gel interface directly as single, lumenized vessels. In turn, non-PMA-stimulated cultures formed an endothelial cell monolayer circumferentially lining the medium channel as visible in Figure 3B irrespective of VEGF-C concentration. In summary, results of our growth factor optimization study revealed a striking morphological similarity of our PMA (100 ng/mL) and VEGF-C (100 ng/mL) co-stimulated vascular networks to blind-ended lymphatic capillaries found *in vivo*.^7^ Consequently, all following experiments are conducted using this co-stimulation protocol to investigate sprouting behavior under dynamic flow conditions.

### Dynamic cultivation initiates sprouting and enhances lymphatic capillary length

Since lymphatic capillaries are highly sensitive to interstitial fluid flow *in vivo*, lymphangiogenesis usually occurs in response to fluid flow irregularities and associated tension variations caused by, e.g., wound healing processes or tumor growth.^43^ To evaluate the impact on LEC sprouting and BEC network formation under unidirectional interstitial fluid flow (e.g., 1 – 3 µm/s) across the hydrogel compartments, the lymphatic sprouting setup (shown in Figure 2A) is used in subsequent experiments. Results shown in Figure 4 clearly demonstrate the impact of dynamic culture conditions on lymphatic sprouting behavior, which led to significantly elevated sprouting activities in both PMA-stimulated and control cultures. Interestingly, dynamic cultivation prompted lymphatic capillary formation even without auxiliary stimulation via PMA and elevated sprouting activity in PMA-stimulated cultures, visible in representative images of capillaries in all conditions in Figure 4A. While PMA-stimulated cultures exhibited a higher number of lymphatic sprouts before fusing into larger capillary structures, unstimulated LEC predominantly formed larger and single capillaries originating from the endothelial monolayer at the hydrogel interface. A detailed quantitative analysis of sprouting activity is shown in Figure 4B, further indicating the generation of elevated sprouting lengths in the presence of dynamic culture conditions. In other words, the presence of unidirectional interstitial flow leads to sprouting length increase by 1.7-fold from 180 µm in static PMA-stimulated cultures to 313 µm in dynamic controls, thus highlighting the beneficial impact of physiologically relevant directional interstitial fluid flow conditions on lymphatic sprouting activity.

**Figure 4:**
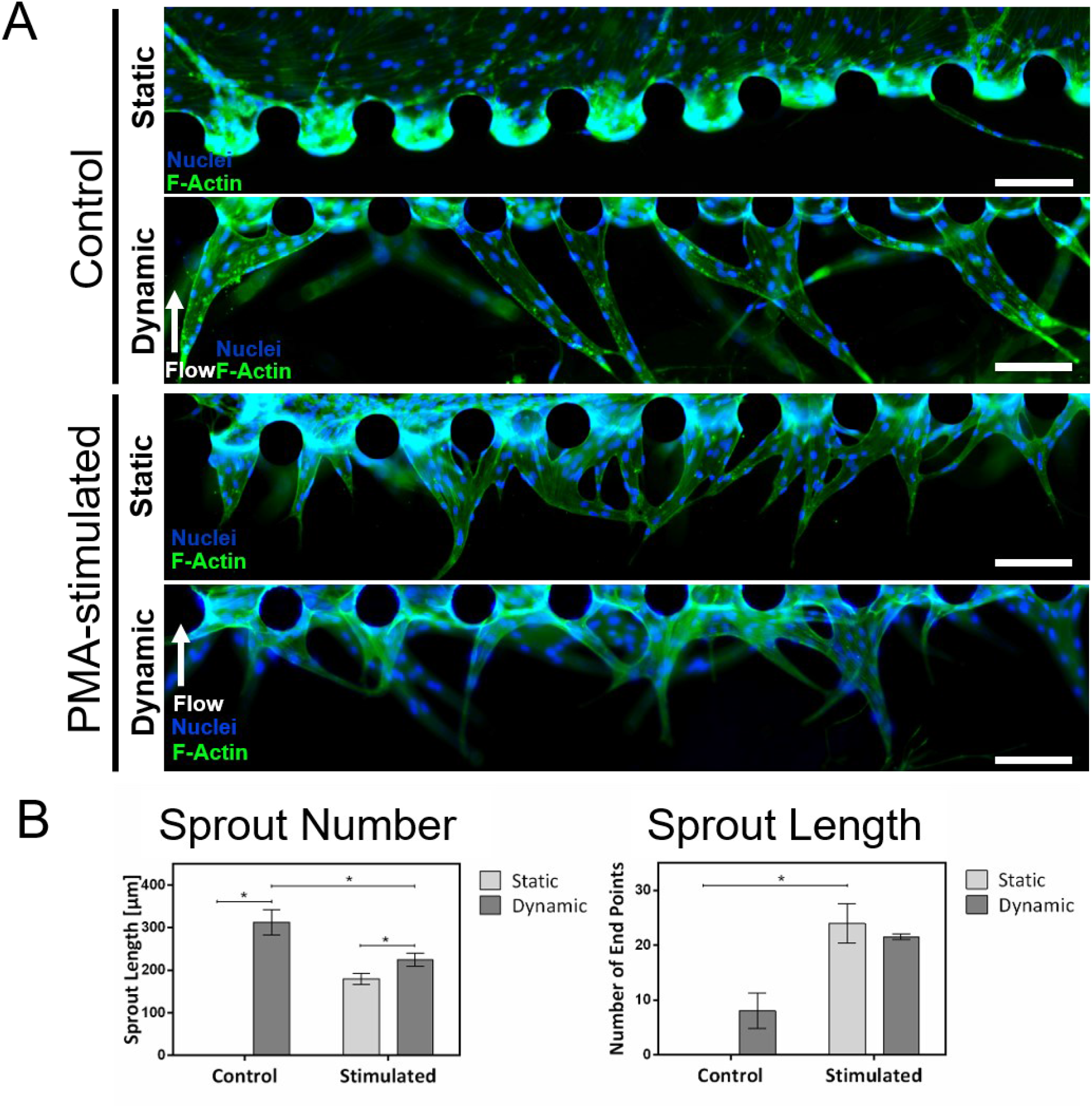
Mechanobiological effect of interstitial fluid flow on LEC sprouting (A) Increase in LEC sprouting activity under dynamic cultivation conditions mimicking interstitial flow. (B) Lymphatic sprout morphology under static and dynamic cultivation conditions under control and in PMA-stimulated cultures. Scale Bar 200 µm. *p<0.05.

Next, we focused on the influence of directional interstitial fluid flow on vascularization behavior of primary blood endothelial cells (BEC) located next to the LEC sprouting compartment and cellular outgrowth from the BEC vascularization chamber towards the lymphatic capillaries, as illustrated in Figure 5. To this end, we conducted a comparative analysis of vascularization parameters in the absence and presence of flow using stimulated and non-stimulated BEC cultures. Figure S4 demonstrates the marginal impact of fluid flow and cell culture stimulation on experimental vascularization parameters. For instance, neither PMA stimulation nor dynamic cultivation significantly impacted vessel area coverages (e.g., 42 % and 45 %), where BEC and ASC co-cultures reliably formed stable networks exhibiting 97 – 114 junctions per network and device. However, average vessel length decreased from an average network length of 10156 µm in statically cultivated control cultures to 5500 µm in dynamically cultivated unstimulated cultures (statistically not relevant). In comparison, PMA-stimulated cultures yielded an increased average vessel length of approximately 1000 µm (e.g., 3230 µm to 4466 µm) under dynamic cultivation. In other words, BEC may react to the constant supply of PMA molecules resulting in increased cell migration and proliferation. Another intriguing effect of the directed interstitial flow conditions was the occurrence of cellular ingrowth from the blood endothelial compartment into the cell-free lymphatic sprouting compartment. Quantitative analyses of cellular protrusion length and nuclei number are shown in Figures 5 and S4, respectively, and demonstrate cell outgrowth predominantly in non-PMA-stimulated cultures, despite supposed migratory stimulation via PMA.

**Figure 5:**
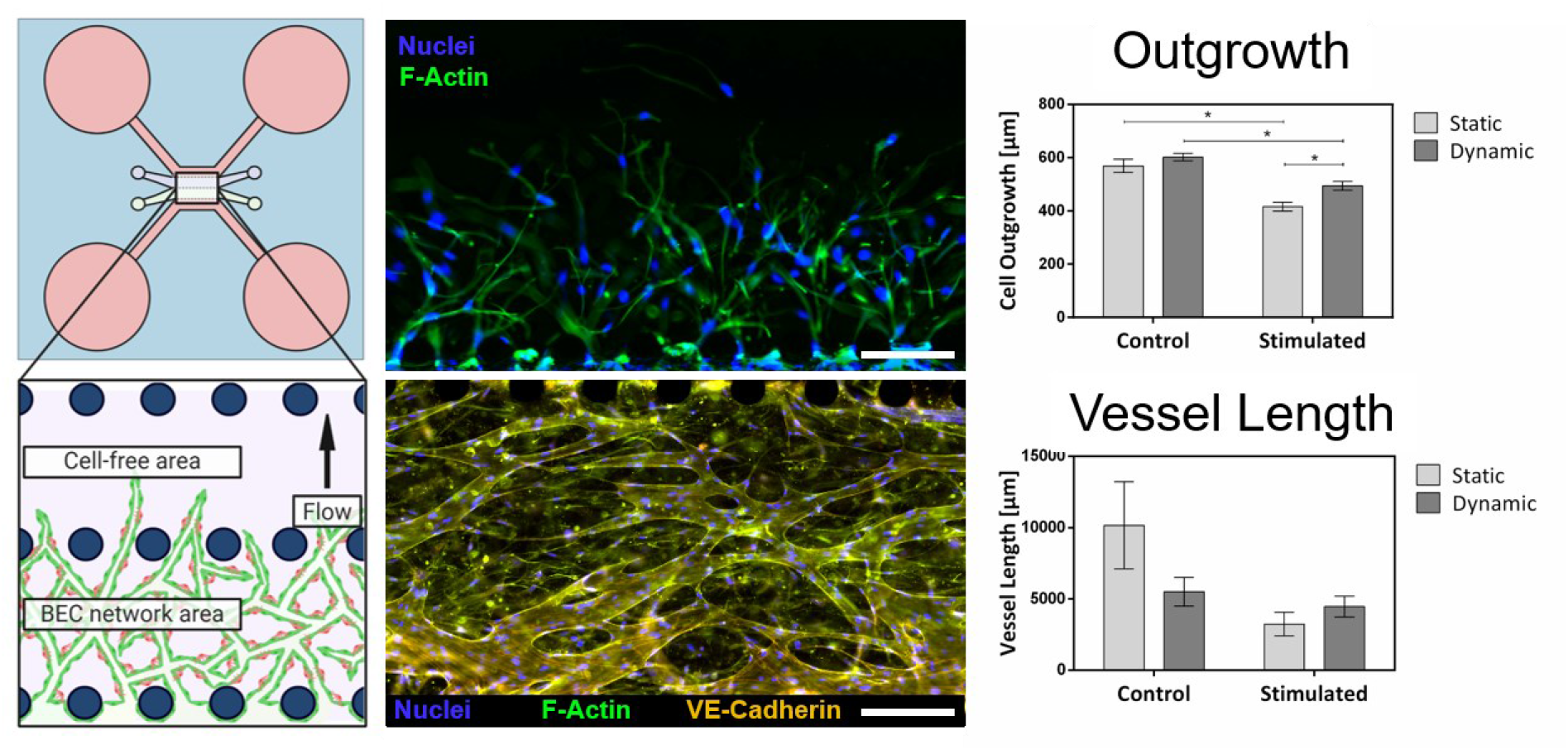
Effect of dynamic cultivation on cell outgrowth from the blood vascular compartment and blood vessel vascularization. Cells in non-PMA stimulated cultures migrate into the gel-free area irrespective of dynamic cultivation, while PMA-stimulated cultures show significantly increased outgrowth under directional fluid flow. BECs form blood vascular networks through vascularization irrespective of flow condition. Scale bar 200 µm. *p<0.05.

In summary, Figure 6 shows the successful establishment of lumenized vessels from blood and lymphatic vascular origin within one cell cultivation unit on-chip. The device supports the formation of vasculatures of both lineages by using our optimized cultivation method and the described lymphatic sprouting setup. Both vasculatures form lumenized vessels expressing endothelial cell-cell junctions and show distinct differences dependent on vasculature type, similar to their *in vivo* morphology. Lymphatic sprouts form thicker blind-ended vessels while blood endothelial cells vascularize the fibrin hydrogel with a thinner, continuous blood vascular network. The microvasculature-on-chip system recreates the interstitial niche by mimicking interstitial fluid flow velocities via hydrostatic pressure-driven flow resulting in enhanced lymphatic sprouting and establishing the blood-lymph interstitial interface within a microfluidic device.

**Figure 6:**
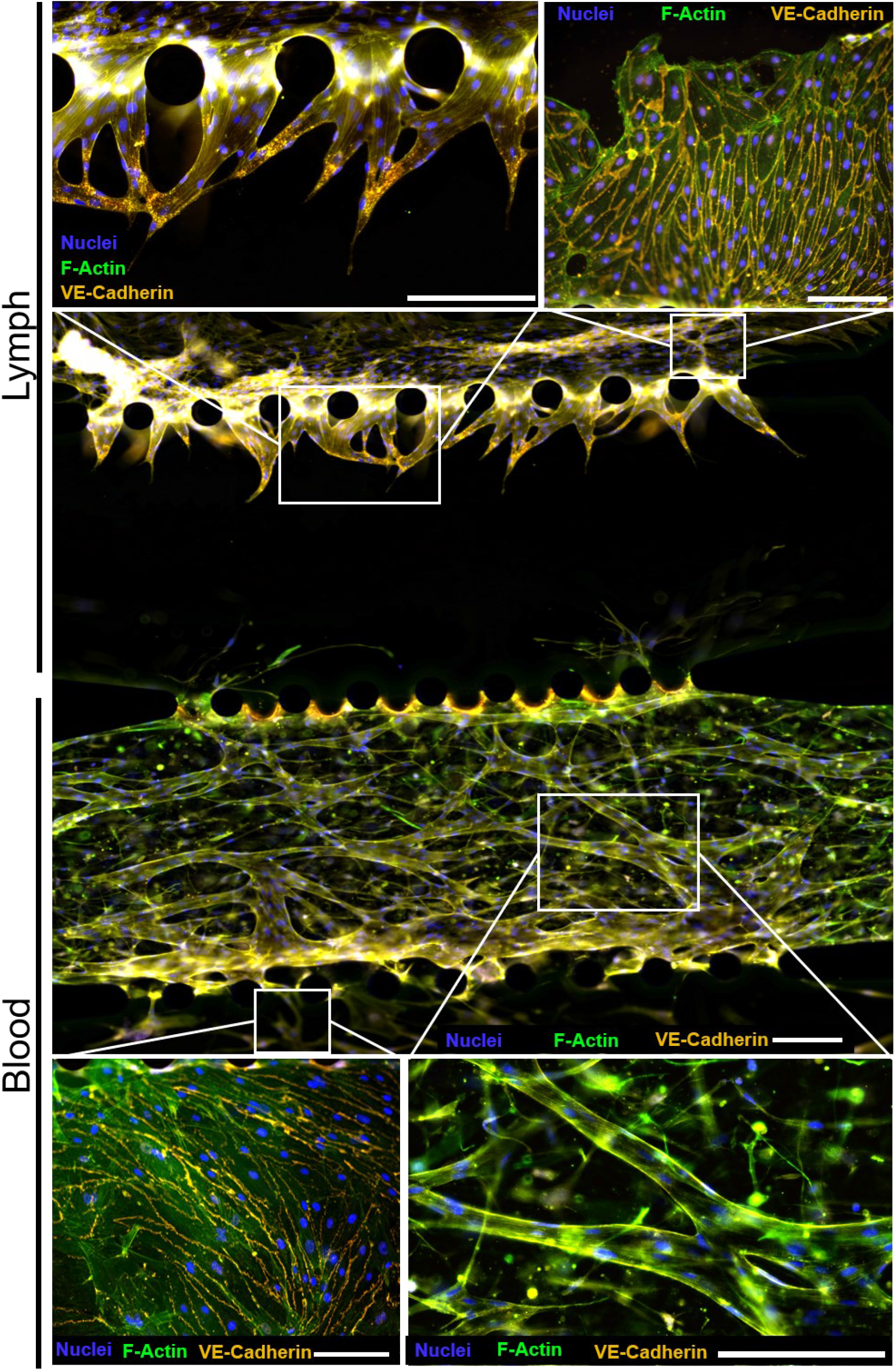
One µLymph unit showing successful lymphatic endothelial sprouting and blood endothelial network formation to generate a blood-lymph capillary bed in a microfluidic device. Lymphatic and blood vasculature exhibit distinct morphology similar to *in vivo* vasculature, form lumenized vessels, and express endothelial cell-cell junctions. Scale bars 200 µm.

## CONCLUSION

In the present work, we investigated our novel microvasculature-on-chip device’s suitability to establish a blood-lymphatic interface i*n vitro*. The device harnesses microfluidic technology’s advantages to tailor the *in vitro* microenvironment by spatial separation of two hydrogel compartments and utilization of hydrostatic pressure-driven flow. Creating hydrostatic pressure via fluid reservoirs allows the generation of interstitial fluid flow of 1 – 3 µm/s to mimic the path of fluid expelled from the blood vasculature traveling through interstitial tissue into the lymphatics. VEGF-C is crucial for the first steps in lymphatic development. Without it, embryonic endothelial cells commit to the lymphatic lineage but do not form sprouts to develop functioning lymphatic vasculature.^41^ We thus utilized our microvessel-on-a-chip platform to evaluate sprouting induction via VEGF-C individually and as a combined effect with PMA, which, too, acts as a potent inducer of endothelial sprouting.^42^ We then demonstrated the impact of mechanobiological stimulated cultivation on spatially separated cultures of blood and lymphatic endothelial cell populations. While vascularization of blood endothelial cells was not affected by dynamic cultivation conditions, dynamic cultivation recreating the interstitial microenvironment enhanced lymphatic endothelial sprout formation. In short, dynamically cultivated lymphatic cultures formed significantly longer lymphatic vessels than their static counterparts. We report the successful formation of both vascular systems within one device, exemplifying the first step towards engineering novel *in vitro* tools to bridge the gap between animal models and conventional monolayer cultures. The establishment of such models will be an essential step to expand our knowledge of the human body in general and the lymphatic system in particular. Our microvasculature-on-chip model will help develop drug delivery strategies into the lymphatic system that can help in advancing modern vaccines and chemotherapeutics.

## CONFLICT OF INTEREST

The authors declare that the research was conducted without commercial or financial relationships that could be construed as a potential conflict of interest.

## AUTHOR CONTRIBUTIONS

BB, WH, and PE designed the study. HW and PS provided expertise in microfabrication and performed photolithography. BB executed experiments and analysis with assistance from SS. CJ, BH, and MH performed computational fluid dynamic simulations. BB, SS, HR, WH, and PE interpreted the data. BB drafted the manuscript with contributions from SS, PE, and CJ. All authors revised and approved the final manuscript.

## FUNDING

This work was partly funded by the European Union’s INTERREG V-A AT-CZ program (ATCZ133).

## ACKNOWLEDGMENTS

The authors acknowledge BioRender.com for facilitating the generation of schematics for the visual representation of the work.

## DATA AVAILABILITY STATEMENT

The raw data supporting this manuscript’s conclusions will be made available by the authors, without undue reservation, to any qualified researcher.

## Supplementary information

**Supplementary Figure 1:**
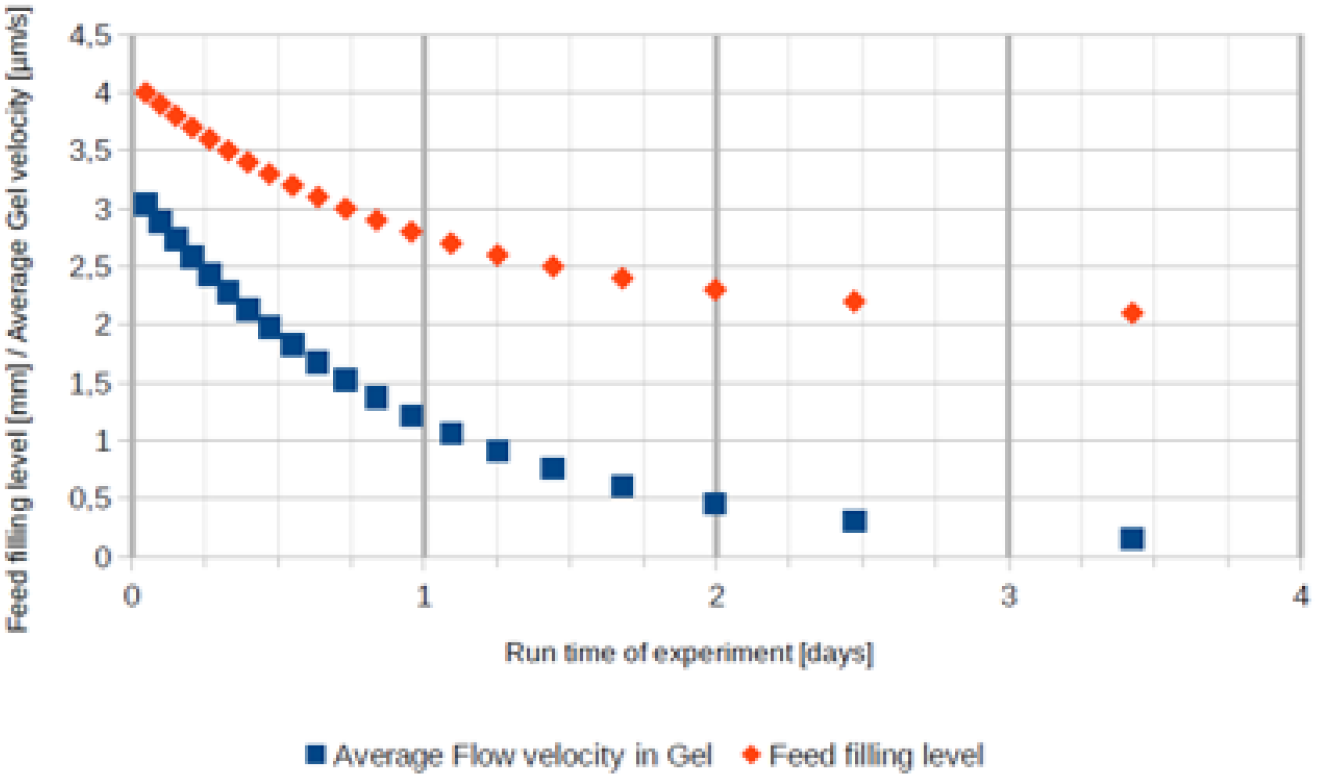
Feed filling level and average velocity in hydrogel over time as determined by CFD simulation.

**Supplementary Figure 2:**
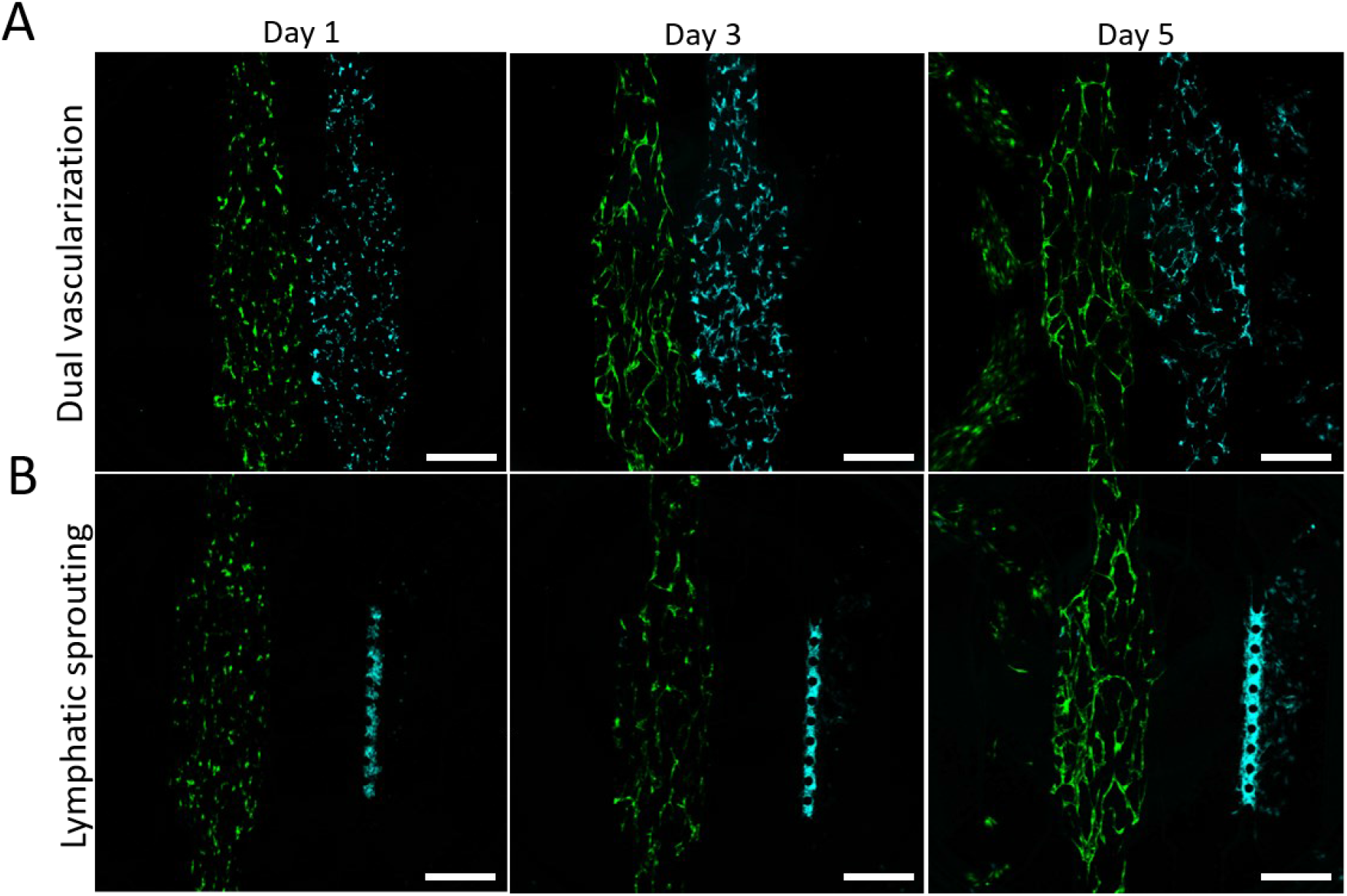
Development of blood-lymphatic interstitial interface over five days of culture time. (A) Dual vascularization setup and (B) Lymphatic sprouting setup. Scale Bar 400 µm.

**Supplementary Figure 3:**
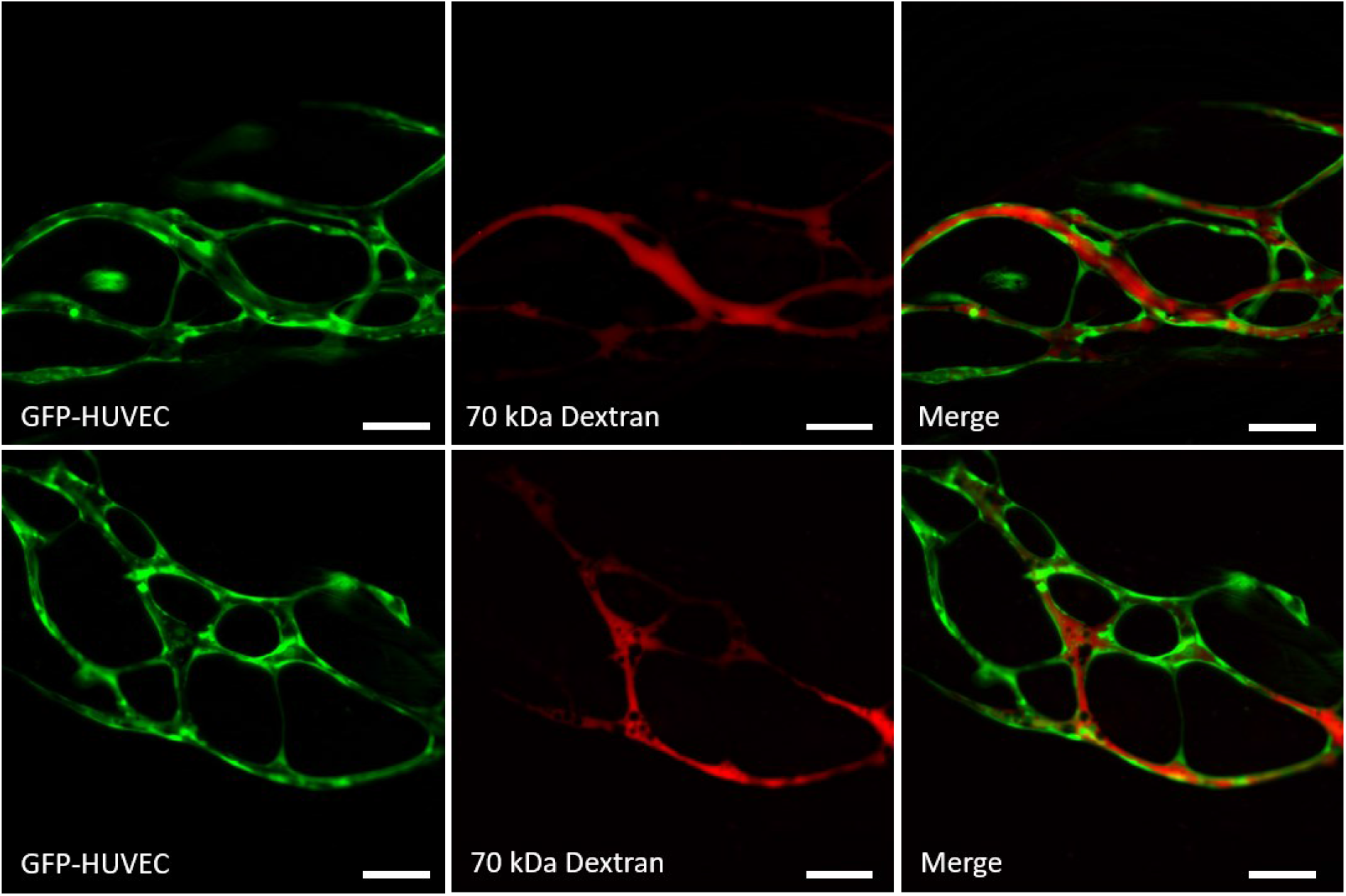
Perfusion of blood vessels (GFP-HUVEC, green) with 70 kDa TRITC-dextran (red) demonstrating lumen formation and vessel functionality. Scale bar 100 µm.

**Supplementary Figure 4:**
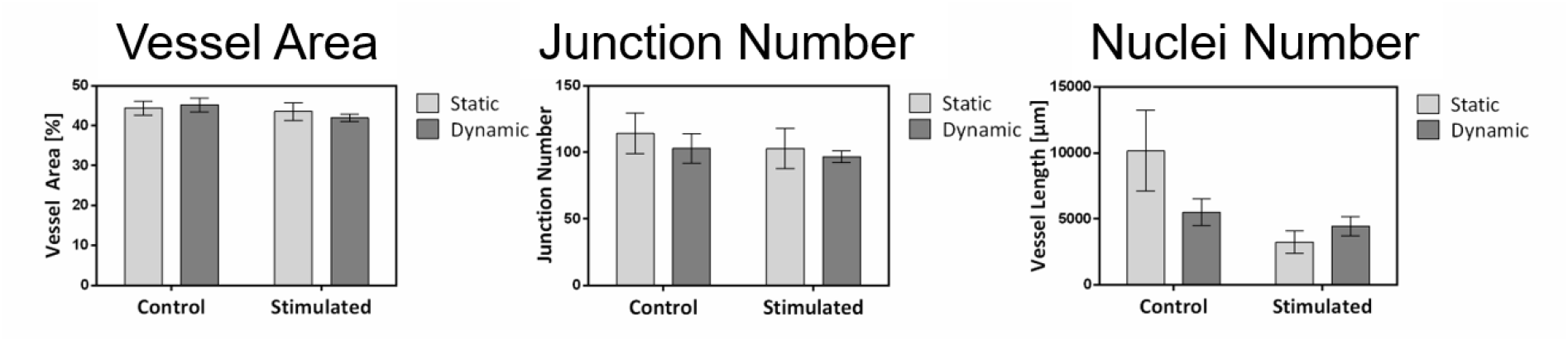
Influence of dynamic cultivation on blood vascular network and nuclei number within the cell-free compartment. The blood vascular network is unaffected by dynamic cultivation showing consistent vessel area coverage and junction number irrespective of cultivation conditions. Cell outgrowth is significantly enhanced in non-PMA-stimulated cultures.

## Notes

### Competing Interest Statement

The authors have declared no competing interest.

